# Detergent-based separation of microbes from marine particles

**DOI:** 10.1101/2025.07.15.664943

**Authors:** Jordan T. Coelho, Lauren Teubner, J. Cameron Thrash

## Abstract

Marine particles, typically composed of organic detritus and cellular debris, harbor microbial communities that are distinct from the planktonic, or free-living, communities in the pelagic ocean. However, without being first separated from the particle and microbial consortia, these microbes are inaccessible to further investigation via single-cell microbiology methods like flow cytometry, cell-sorting, and dilution-based isolation. To confront this obstacle, we compared the dissociative effects of two commonly used detergents, Tween20 and Tween80, on particle-associated marine microbial communities. The ability of Tween treatments to liberate cells from particles, and to maintain cell integrity, was quantified by flow cytometry from multiple communities across seasons and locations. Both Tween20 and Tween80, at 185 RPM shaking, gently dissociated microbes from their particles, causing very little cell mortality. Additionally, Tween80 liberated a greater number of particle-associated cells into the free-living fraction. We also analyzed the effects of Tween treatments on the microbial community composition for one of these collections via 16S rRNA gene amplicon sequencing of the particle-associated and free-living fractions relative to unamended controls. Tween20 and Tween80 were both effective for microbial dissolution from particles, however Tween80 treatments displayed greater uniformity in the dissociated communities and significantly enriched for the most abundant particle-associated members. Together these data indicate that Tween80 was most effective at gently dissociating particle-associated cells.

**Importance:** Microbes that reside on marine particulate organic matter are vital facets of marine biogeochemistry. As they degrade the particle on which they reside, the resulting concentrated region of activity influences surrounding biogeochemistry and redox gradients, making particle-associated microbes significant to overall marine ecology. To understand single-cell activities amidst the microbial assemblage on the particle, cells must first be removed from the substrate for downstream analyses. Methods for microbial dissociation from solid surfaces or sediment communities have been described, however, analogous methods for more ephemeral particles, that also maintain cell viability and preserve DNA for next-generation sequencing, are understudied. Here we optimized a method that leveraged detergents to dissociate microbes from marine particles. We evaluated effectiveness through filter size-fractionation, flow cytometry, and community composition analyses, and provide recommendations to gently and effectively remove microbes from marine particles.

## Introduction

Marine particles result from a wide range of biotic and abiotic sources [1] and are largely comprised of aggregates of cellular debris, fecal pellets, and organic detritus [2]. These particles are broadly defined as particulate organic matter (POM), and due to their density in nutrients and carbon, represent hot-spots of microbial activity where microbes transiently or permanently reside [3–5]. Particle-associated (PA) microbes form complex communities that coordinate their efforts to degrade the POM [6, 7], and often the PA microbial assemblage reflects the array of metabolic niches present within the particle [8–10]. PA communities are taxonomically distinct from planktonic/free-living (FL) communities, and can be influenced by differing environmental conditions, with genomic evidence suggesting strikingly different physiologies between the two communities [11–14]. Thus, PA communities occupy separate ecological niches than those of FL communities. However, PA communities are generally inaccessible to single-cell microbiology methods [15] including flow cytometry and fluorescence-activated cell sorting (FACS), dilution-to-extinction cultivation methods, and single-cell genomics, all of which have greatly contributed to our understanding of the ecophysiology of many important microbial lineages and ecological processes [16–22]. For example, due to PA microbes living in dense consortia inside or on the surface of particles, a flow cytometer cannot distinguish individual cells, and particles may obstruct the fluidics lines of the instrument. Other single-cell methods face similar challenges. Thus, dissociation of PA microbes from their particles makes them available for a wider array of methods to investigate their ecophysiologies.

Many methods have been developed to separate microbes from surfaces. An effective approach for detaching particle-associated and sedimented-associated microbes involved 10% (v/v) methanol and sonication [23]. Although the methanol disruption approach has been applied across marine particles [24], ruminal digesta solids [25], and deep-sea sediments [26], 10% (v/v) methanol requires a large chemical addition to the sample and is toxic to microbial cells [27], thus inhibiting growth and propagation of the dissociated cells. Other methods using low pH, formaldehyde, tertiary butanol, and methylcellulose [25, 28] have similarly toxic effects on microbial cells, and some also prevent downstream DNA sequencing [29]. Density centrifugation is another method that has been successful in detaching sediment-associated cells, however the protocol is lengthy, the reagents are expensive, and the multiple centrifugation and supernatant removal steps can lead to cell losses [26, 30].

Detergents are a popular choice for dissociating microbes from sediments [31], soils [32], surfaces in the food industry [33, 34], and atmospheric particles [35] because they disrupt the linkages between the microbe and the particle[35]. Among the commonly used detergents are Pyrophosphate and Tween, representing ionic and nonionic detergents, respectively. Ionic detergents contain a charged polar head group that elicits a strong destabilizing effect, making them ideal constituents in cell lysis buffers [36, 37]. However if the goal is to maintain cell membrane integrity and viability, ionic detergents are not the optimal choice due to their high affinity for proteins [38] and lipid membranes [39]. Nonionic detergents, with an uncharged polar head group, are a gentler option for disrupting linkages between microbes and particles. Furthermore, the ionic strength of the surrounding matrix has little to no effect on the efficacy of nonionic detergents [40], making them a suitable choice for marine applications.

The Tween family of nonionic detergents are frequently used for stabilizing enzymes [41], isolating proteins [42], and dilute concentrations have previously been used for dissociating microbes from sediments and atmospheric particles for downstream flow cytometry applications [30, 31, 35]. Tween20 and Tween80 are the two most commonly used Tween detergents, however, their efficacy has not yet been proven for the removal of microbes from marine particles. The same concentrations and variations of these detergents that have been previously proven effective on sediment [30, 31] or atmospheric [35] microbial communities, among others, may not have the same effect on marine PA communities because of the fragile and ephemeral nature of marine particles, the distinct and dynamic communities, processes, and interactions occurring on them [6, 43, 44], and the multiple mechanisms that microbes use for attachment [45].

We tested the efficacy of Tween20 and Tween80 for liberating cells from marine particles. We quantified the effectiveness of Tween treatments in samples from different seasons and locations using flow cytometry to enumerate the number of viable cells dissociating from the PA fraction into the FL fraction and analyzed the PA and FL microbial community composition via 16S rRNA gene amplicons after Tween treatments relative to unamended controls to determine which organisms were being affected by the treatment. Tween80 treatments consistently enriched for the most abundant particle-associated taxa, whereas Tween20 treatments were more variable and enriched for rarer particle-associated taxa, corroborating flow cytometry observations that low concentrations of Tween80 were the most effective for dissociating PA microbes from marine particles.

## Results and Discussion

### Experimental design

We developed an experimental plan based on a common sequential filtration procedure to separate particle-associated (PA) from free-living (FL) marine microorganisms. We chose 2.7 µm as a cutoff to define PA communities and 0.22 µm to capture the majority of remaining cells (**Fig. 1**). Our experimental logic was that an effective dissociative treatment that liberated cells from particles would result in a greater proportion and diversity of microbes passing through the 2.7 µm filter than without such a treatment. Our treatments consisted of the commonly used detergents, Tween20 and Tween80, that we tested at four concentrations spanning previous ranges used for liberation of cells from sediments (0.001%, 0.005%, 0.01%, and 0.1% (v/v)), along with controls that did not receive Tween. We then used flow cytometry to measure changes in bulk cell numbers, and viable cells, across multiple treatment types and samples (**Fig. 1A**) and community composition analyses (**Fig. 1B**) to compare the difference in microbial cells passing through the 2.7 µm filter.

**Figure 1.**
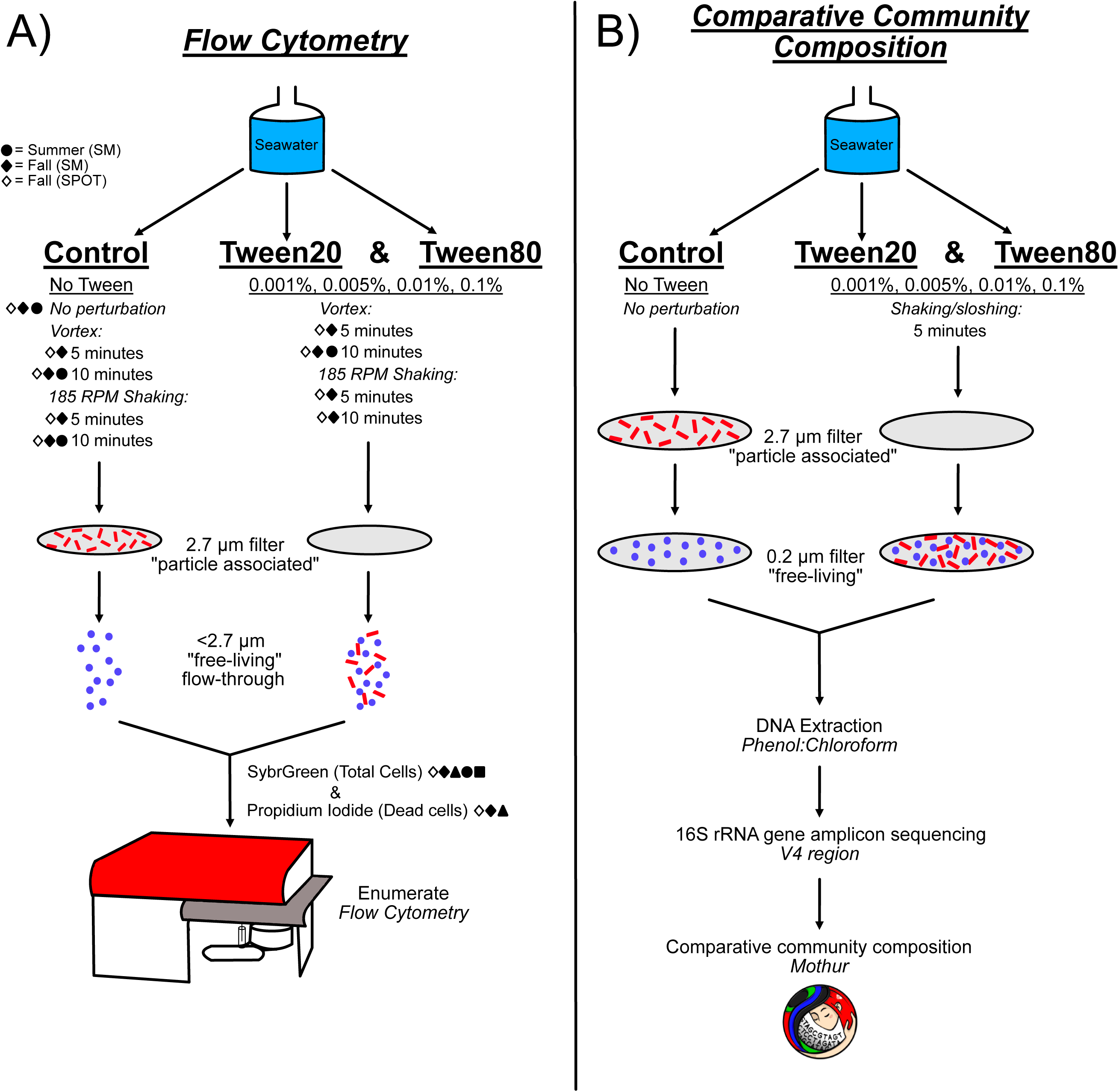
Experimental design and workflow. Experimental conditions are described with underlined text, and the perturbation method is described by italicized text with the duration of perturbation listed below. The filter size-fractionation of the particle-associated community (red) and the free-living community (blue) is depicted by the grey discs that represent the filters, with the particle-associated community becoming enriched in the planktonic size fraction after Tween treatments. A) Experimental design and workflow for the verification of microbial dissociation from particles via flow cytometry. Not every seawater collection received the same perturbation methods and durations, therefore the shapes indicate the season and location from which seawater was collected for experimentation: Winter, Santa Monica (◾), Spring, Santa Monica (⚫), Summer, Santa Monica (▴), Fall, Santa Monica (◆), Fall, SPOT (◇). B) Experimental design and workflow for the verification microbial dissociation from particles via 16S rRNA gene community composition analysis.

### Cellular dissociation and viability

We evaluated the dissociation efficiency of Tween treatments via flow cytometry at two different sampling sites: Santa Monica Bay, to represent a coastal system, and the San Pedro Ocean Time series station (SPOT), to represent an open ocean system. We quantified total and compromised FL cells after five or ten minutes of shaking at 185 RPM relative to no-Tween shaken controls and no treatment controls (**Fig. 2**). In our Fall 2023 Santa Monica Bay experiment, we observed the greatest liberation of cells with 10 minutes of shaking in Tween80 0.005%, 0.01%, and 0.1%, yielding median percent increases in cell density from the control of 69.3%, 75.6%, and 70.7%, respectively (**Fig. 2A**). Cell mortality was similar across all Tween/shaking treatments at below 2% of the total cell count. Controls showed roughly 1% cell mortality. Our Spring 2024 SPOT experiment had different absolute numbers, but the trends corroborated our previous experiment. Tween20 and Tween80 had more similar results at 5 minutes of shaking, but at 10 minutes of shaking Tween80 yielded slightly greater cell dissociation, based on the percent cell increases relative to controls (**Fig. 2B**). We observed the highest dissociation with 10 min shaking in Tween80 0.001%, 0.005%, and 0.01%, with median percent increases in cell density from the control of 52.7%, 50.3%, and 54.6%. Cell mortality was similar between both Tween20 and Tween80 and represented less than 4.5% of the total cell count, compared to ∼2-3% mortality in the controls. The higher mortality observed in the SPOT samples compared to Santa Monica Bay could have resulted from differences in sample collection and processing times (see Materials and Methods), however, both experiments indicated that 10 min shaking with Tween80 at 0.005-0.01% dissociated the most PA cells into the FL fraction.

**Figure 2.**
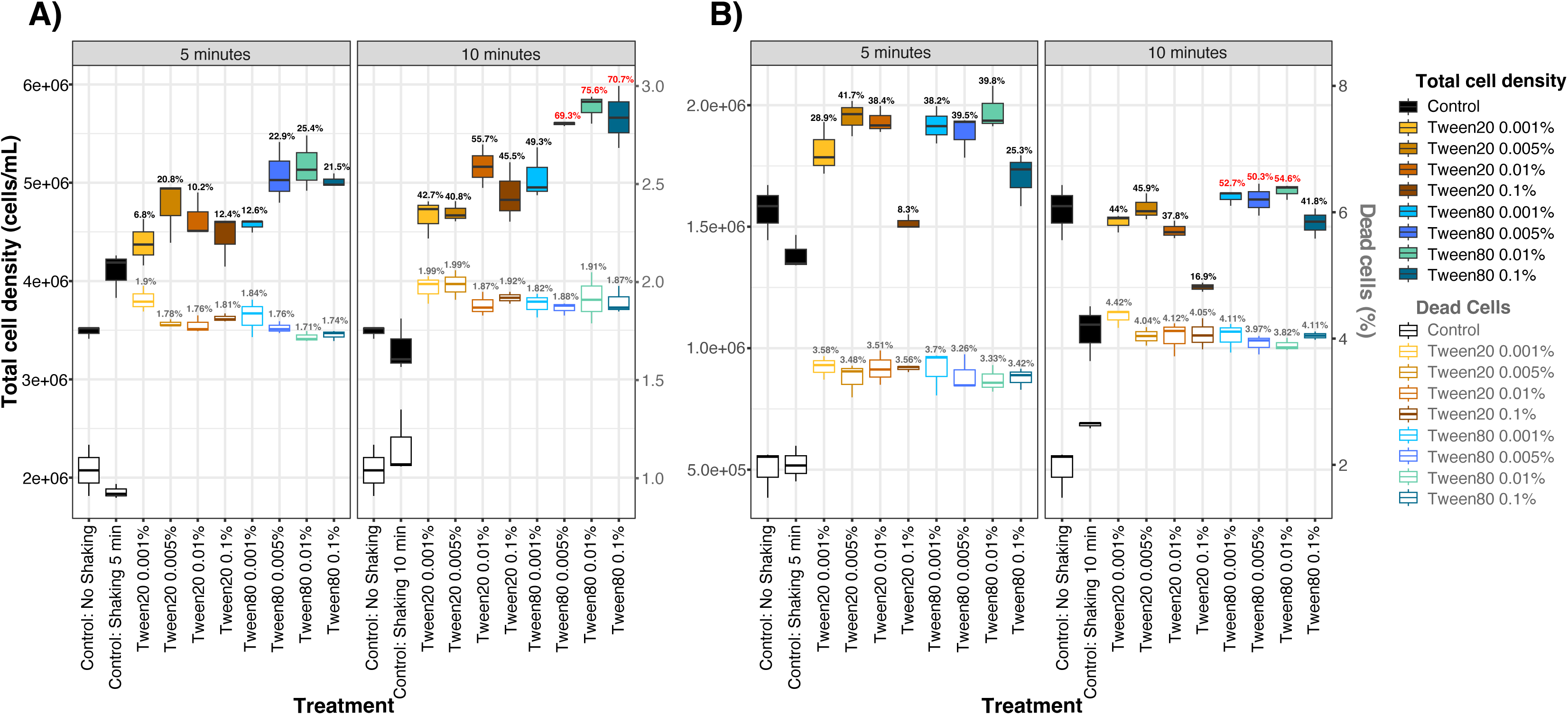
Microbial dissociation from marine particles evaluated by flow cytometry, for the Santa Monica Bay, (A) and SPOT (B). X-axes indicate treatments. The left y-axis displays the total cell density because of the SYBR-Green fluorescent stain (black text), and the right y-axis displays the cell density of the compromised cells within the sample because of the Propidium Iodide fluorescent stain (grey text). The data are separated by the duration of perturbation, either 5 or 10 minutes of 185 RPM shaking. The boxplots show the lower and upper quartiles with the horizontal line indicating the median and the whiskers showing the minimum and maximum values. The percentages above the boxplots indicate the average percent increase in cell density, or percent cell death, relative to the controlthat received physical perturbation. The top three most successful Tween conditions that dissociated the most particle-associated cells are colored in red.

Variations to the perturbation type and duration confirmed Tween80 and ten minutes of 185 RPM shaking to be the most consistent treatments. Experiments using Santa Monica Bay surface water across multiple seasons showed that vortexing introduced more variability in dissociation frequency, and in some cases drastically increased cell mortality, compared with 185 RPM shaking (**Fig. S1**). Direct comparison of 10 min shaking vs vortexing using a summer surface sample showed comparable results (**Fig. S1A**). However, when comparing 5 vs. 10 min vortexing with a fall sample, we found very high cell mortality in the 10 min treatment, exceeding 2-3% of the untreated controls (**Fig. S1B**). Comparing cell mortality after 10 min vortexing in the summer (**Fig. S1A**) vs the fall (**Fig. S1B**) suggested that vortexing differentially affected cell mortality depending on the presumed variation in microbial community composition between seasons. Shaking at 185 RPM provided comparable or higher dissociation of cells than vortexing, and with more consistent outcomes in cell mortality across summer and fall seasons (**Fig. 1, S2A**). Collectively, our experiments indicated that Tween80 with gentle shaking for 10 min reliably dissociated PA cells from a variety of different sample types, and frequently at higher numbers than Tween20.

### Community effects

To evaluate the effect of the dissociation treatments on the microbial community and to identify which taxa were liberated from particles, we used whole community 16S rRNA gene amplicon sequencing of communities obtained from surface water, treated with the same suite of Tween conditions and gentle manual shaking of carboys (roughly equivalent to 185 RPM shaking) described above. As expected based on many prior observations [12, 46–49], the PA and FL communities were distinct as evaluated by principal-coordinate analysis, with PCo1 separating the PA and FL communities and explaining 78.18% of variance (*R^2^*=0.96) (**Fig. 3**). PCo2, which appeared related to the Tween treatments, explained only 4.46% of variance (*R^2^*=0.98). In both the PA and FL communities, the community structure diverged away from the tightly clustered control communities after Tween treatments. We also observed a slight movement in the FL communities along PCo1 after treatment, suggesting that some liberated PA taxa were affecting the composition of the FL community. Replicates of PA communities were highly divergent and did not form consistent clusters based on Tween treatments, suggesting a high degree of heterogeneity in the dissociative results. To quantify the degree of heterogeneity across treatments, we examined the percentage of operational taxonomic units (OTUs) that were shared or unique among the replicates in each of the treatments (**Fig. S2**). An average of 43% of OTUs were unique to a single replicate across both the PA and the FL communities, indicating considerable variation in the microbial communities collected across filters and treatments, and corroborating the observed variance in PCo2 across treatments. We detected an average of 37% of the OTUs across all three replicates in both the control PA and control FL treatments. Therefore, we separately defined PA-OTUs and FL-OTUs as those that were detected across all three replicates in control 2.7 µm and 0.22 µm filters, respectively, and then tracked changes of these “core” OTUs to gain a meaningful understanding of how their relative abundances changed in response to Tween treatments.

**Figure 3.**
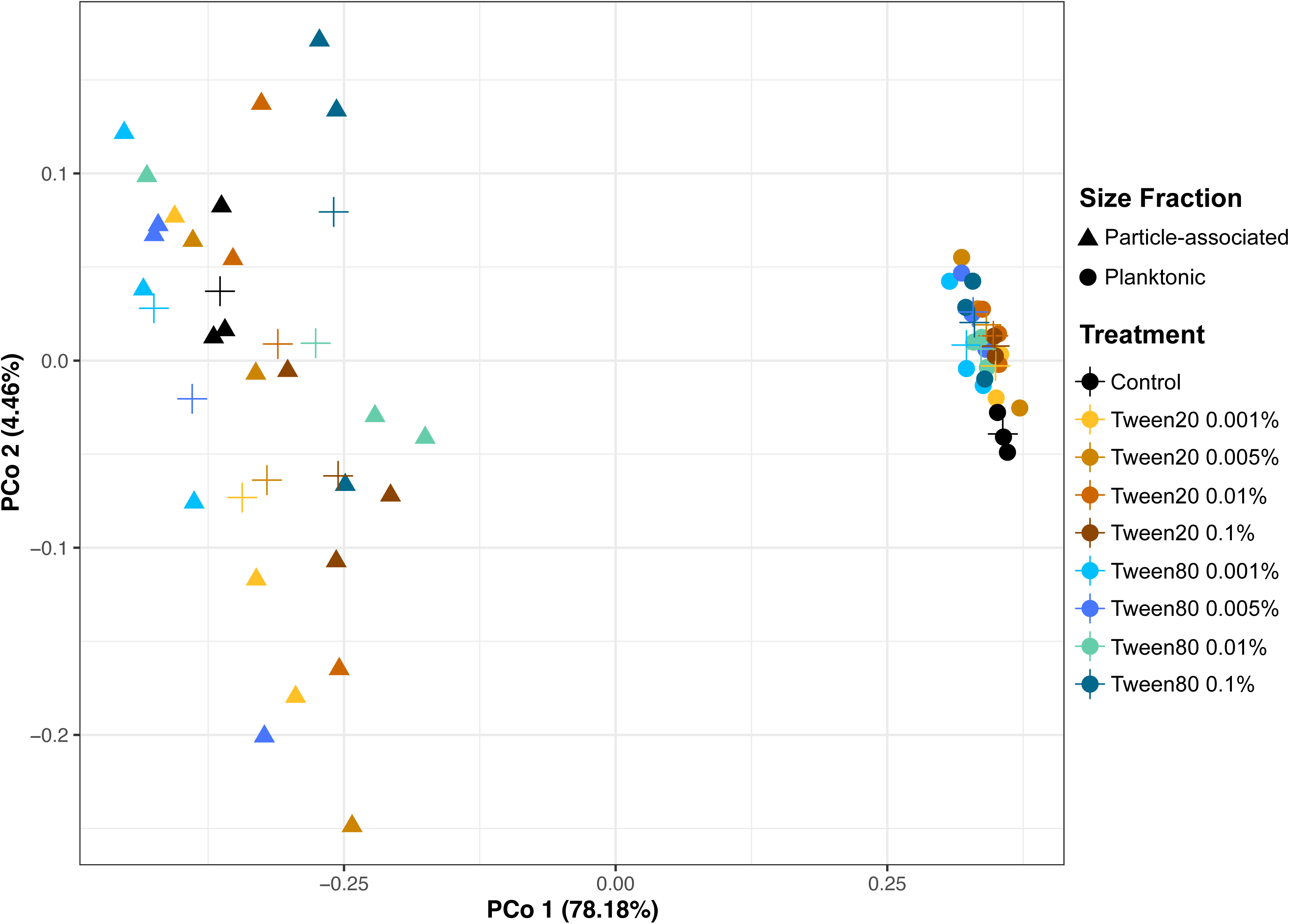
Principal Coordinate Analysis of the particle-associated and free-living communities in response to Tween treatments. Shapes on the plot indicate either particle-associated (▲) or free-living (⚫) communities, and the color of the shapes correspond to the experimental condition or control, according to the key. The crosses indicate the centroid points for each treatment.

### OTU Specificity

Successful microbial dissociation from particles would result in these PA-OTUs becoming enriched in the FL fraction (0.22µm - 2.7µm). We calculated the fold change in PA-OTU relative abundance in the FL fraction after Tween treatments and identified 316 PA-OTUs that increased by more than 1-fold in at least one Tween treatments (**Table S1**). Using principal coordinate analysis of these 316 PA-OTUs, we found distinct communities were liberated from Tween80 and Tween20 treatments, with the Tween20 0.1% treatment diverging from all other treatments (**Fig. 4**). OTUs liberated from Tween80 treatments showed less variation in community composition compared to those liberated from Tween20 treatments overall, implying a more consistent dissociation effect from Tween80. The first two principal coordinates, PCo1 and PCo2, collectively explained only 40.2% of the variance in the community composition, which suggests that the differences in PA-OTU composition between Tween treatments was complex and the relationships were not easily reduced to the principal coordinates. Nevertheless, these results demonstrate that the choice of Tween treatment affects not only the magnitude of cellular dissociation (**Fig. 1**), but also the consistency of the enriched microbial community.

**Figure 4.**
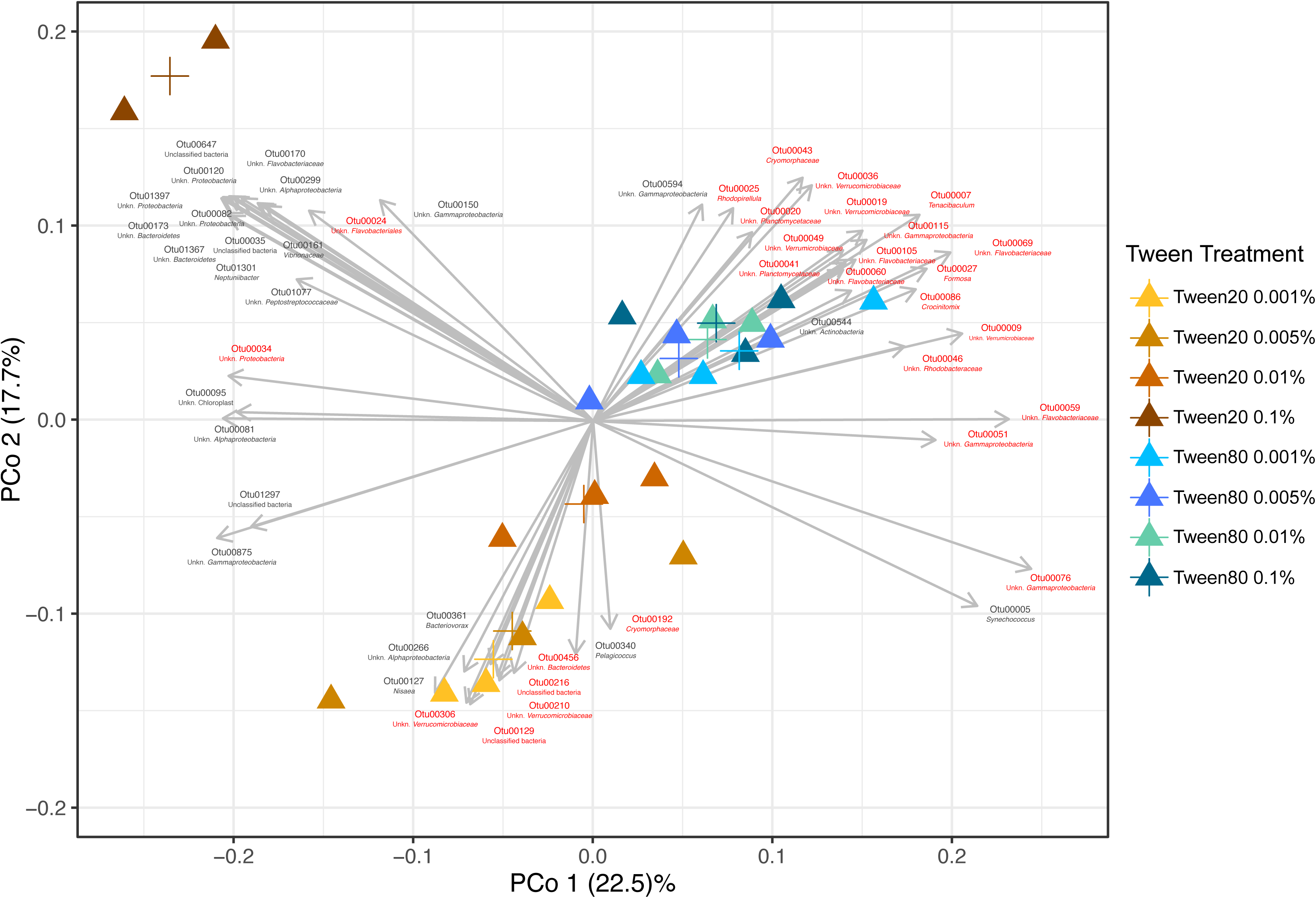
Principal Coordinate Analysis of the particle-associated communities that increased in the free-living fraction in response to Tween treatments (P value < 0.01). The colors of the particle-associated (▴) shapes correspond to the Tween treatment, as indicated by the key. The crosses indicate the centroid points for each treatment. Vectors represent increased PA-OTUs with significant relationships (P value < 0.01) to the ordination axes, which are driving the differences in PA-OTU community composition between Tween treatments. Red vector labels represent E-PA-OTUs that increased significantly (P value < 0.05) in the FL fraction in response to at least one Tween treatment relative to controls (Figure 6, Table S3).

We observed that different PA-OTUs drove the separation between Tween20 and Tween80 communities, and that these differences were not associated with taxonomy (**Fig. 4, Table S1**). For example, PA-OTUs that were highly correlated with Tween80 treatments included members from the phyla *Proteobacteria, Bacteroidetes, Verrucomicrobia*, *Actinobacteria*, and *Planctomycetes*, but we also observed this broad taxonomic representation in PA-OTUs driving variation across Tween20 treatments (**Fig. 4**). Instead of a taxonomic effect, we observed that OTU abundance was associated with the differences in enrichment by either Tween20 or Tween80. OTU abundance corresponds to OTU number (00001 being greatest), and many of the most abundant PA-OTUs corresponded with the Tween80 treatments in the principal coordinate analysis, whereas the PA-OTUs correlated with the Tween20 treatments were less abundant or rare taxa, except for two OTUs associated with the Tween20 0.1% treatment (**Figs. 4, S3**). When using a 0.1 % relative abundance cutoff to define OTUs as “abundant”, of the 65 PA-OTUs at or above that cutoff, 21 of them were correlated with Tween80 treatments and only three with Tween20 treatments (**Figs. 5, S3**). Additionally, we defined enriched PA-OTUS (E-PA-OTUs) as those that significantly increased in at least one Tween treatment relative to the FL control n = 85, *P* value ≤ 0.05, **Fig. 6**, **Table S3**). The majority of PA-OTUs that correlated with Tween80 treatments in the principal coordinate analysis consisted of E-PA-OTUs, compared to Tween20 treatments that corresponded to relatively few E-PA-OTUs **Figs. 4, S3**, **Table S3**). Altogether, these results indicated that Tween80 treatments enriched for taxa that best represented the PA microbial fraction.

**Figure 5.**
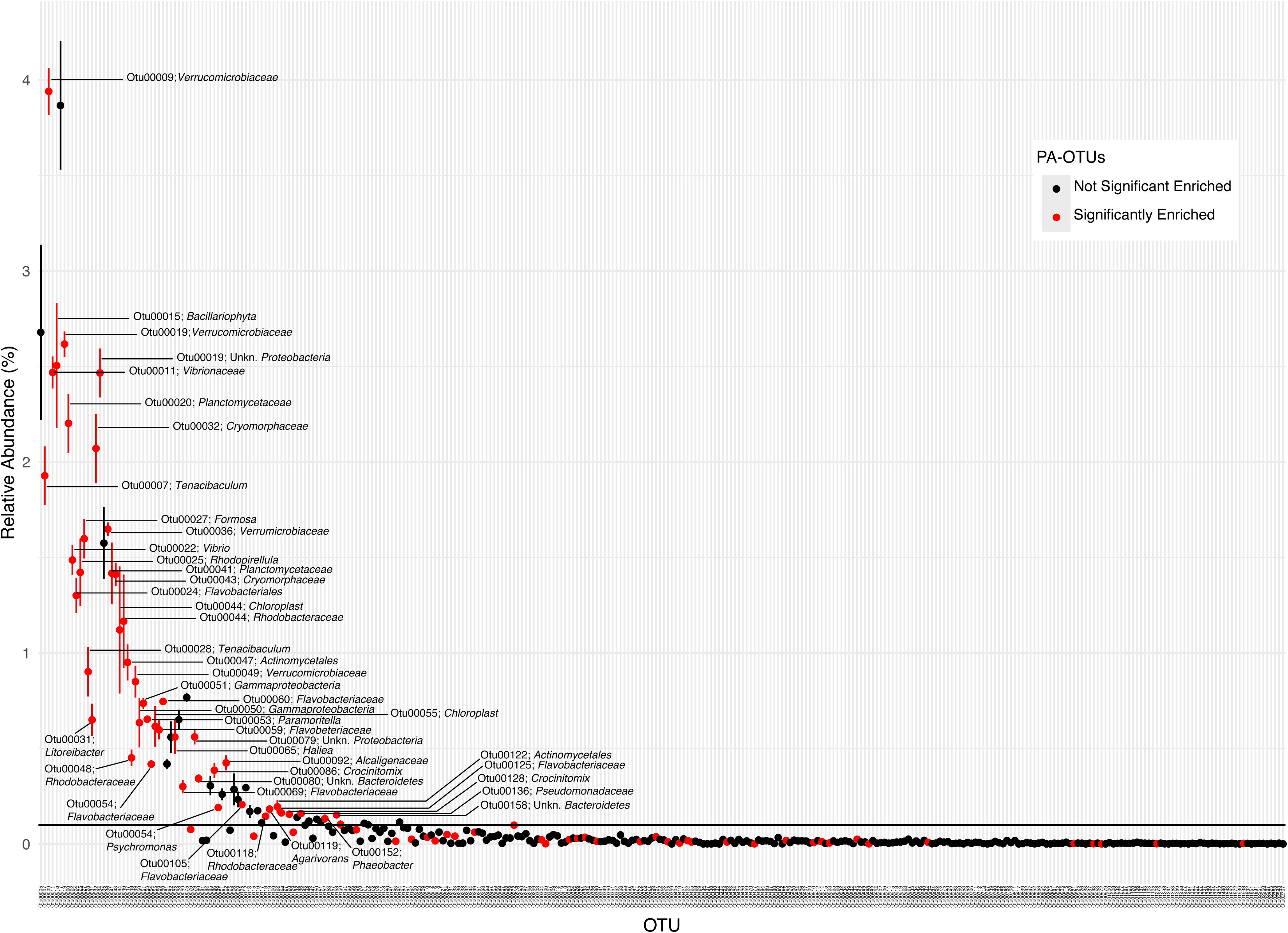
Rank abundance curve of the increased PA-OTUs and classification as abundant or rare. The x-axis shows individual OTUs in rank-order, and the y-axis indicates their percent relative abundance. The black horizontal line shows a relative abundance of 0.1%. An OTU was considered “abundant” if it had a relative abundance > 0.1% in the control PA treatment. Datapoints and error bars represent the mean and variation in fold-change in relative abundance across triplicates. Red data points indicate E-PA-OTUs that were significantly enriched in the FL-fraction in response to at least one Tween treatment (Figure 6, Table S3).

**Figure 6.**
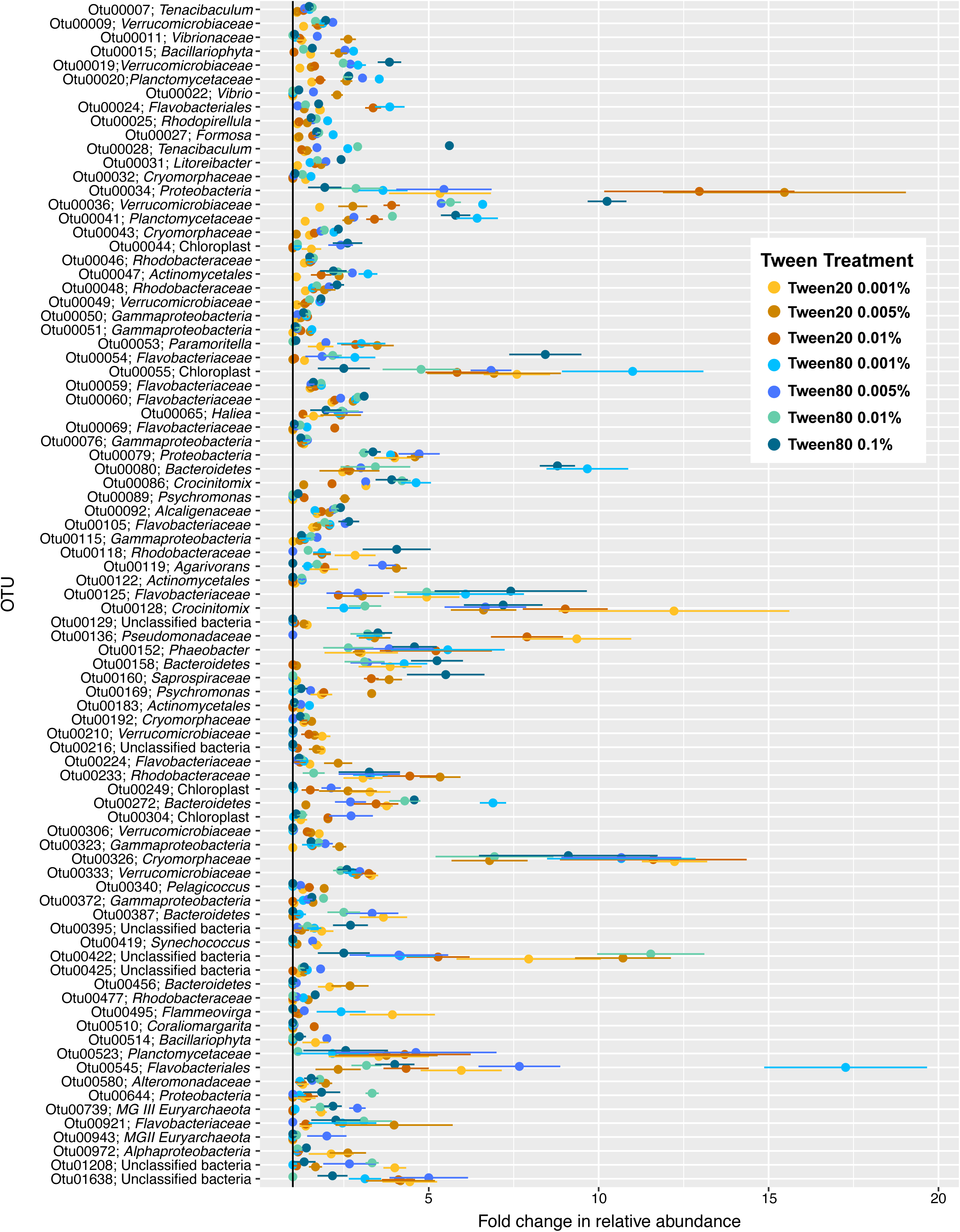
E-PA-OTU enrichment by Tween treatment. The E-PA-OTUs that significantly increased in at least one Tween treatment (Table S3) are organized by OTU number (most overall relative abundance to least). The x-axis indicates the fold change in relative abundance, and the y-axis shows OTUs with their taxonomy. The colors of the data points correspond to the Tween treatment, as indicated by the key. Datapoints and error bars represent the mean and variation in fold-change in relative abundance across triplicates.

### Abundant vs. Rare PA-OTU enrichment

The fact that more abundant E-PA-OTUs were correlated with Tween80 treatments than with Tween20 (**Figs. 4, S4**) did not mean that these were never enriched by Tween20. Rather, Tween80 treatments enriched these OTUs more than Tween20 (**Fig. 6**), similarly to how Tween80 liberated more viable cells overall (**Figs. 2, S1**). For example, one of the most abundant E-PA-OTUs, Otu00009 (member of *Verrucomicrobiaceae,* a commonly detected PA group [8, 50–55]), was enriched across all Tween treatments, but the greatest change in relative abundance was in the Tween 80 treatments, particularly Tween80 0.005%, where its abundance more than doubled (**Figs. 5, 6, S4**, *P* value = 0.04). Other abundant E-PA-OTUs, Otu00019 and Otu00028, members of the *Verrucomicrobiaceae* and *Tenacibaculum* within the *Flavobacteriaceae*, respectively (also typical PA community groups [8, 56]), were slightly enriched in Tween20 treatments, but were much more enriched among Tween 80 treatments (**Fig. 6**). Upon visual (**Fig. 6**) and statistical inspection (**Table S3**), the majority of these taxa were more highly enriched in Tween80 treatments compared to Tween20 treatments (**Fig. 6**), particularly the taxa with numerical OTU identifiers less than Otu00050 (**Fig. S5**), thus corroborating PCoA results (**Figs. 4, S3**). Additionally, all of these abundant E-PA-OTUs matched taxa that have been previously identified in PA communities [3, 8, 50, 57–60], confirming the taxonomic enrichment trends that we detected.

Although the majority of abundant E-PA-OTUs became enriched in the Tween80 treatments, there were a few that were correlated with, and better enriched by the Tween20 treatments instead. Two of these -Otu00024 and Otu00034, unclassified members of *Flavobacteriales* and *Proteobacteria*, respectively - were strongly correlated with Tween20 treatments (**Figs. 4, S3**), primarily driven by the highly divergent Tween20 0.1% treatment (**Figs. 4, S5**). Tween20 at 0.1% caused average fold change increases of 11 and 27 for Otu00024 and Otu0034, respectively, almost double the fold change increases for these OTUs across all other treatments (**Fig S5**). Otu00136 and Otu00169, members of *Pseudomonadaceae* and *Psychromonas* of the *Gammaproteobacteria*, were also correlated with Tween20 treatments (**Fig. S3**), but were predominantly driven by Tween20 0.001, 0.005, and 0.01% treatments (**Fig. 6**). Otu00136 was most highly enriched by Tween20 0.001 and 0.01%, increased by approximately 6-fold and 9-fold in each treatment, respectively. Whereas Otu00169 increased approximately 3-fold in the Tween20 0.005%. The enrichment of these OTUs in the Tween20 treatments may indicate that the way in which they adhere to particles, or the particulate matrix on which these organisms reside itself, was particularly responsive to Tween20 dissolution.

Rare E-PA-OTUs included numerical OTU identifiers greater than Otu00184 (**Fig. 5, Table S2**), and upon visual (**Fig. 6**) and statistical inspection (**Table S3**), the majority of these taxa were more highly enriched in Tween20 than Tween80 treatments (**Figs. 6, S5**). Otu00192, a member of *Cryomorphaceae* (a commonly detected lineage within PA communities [56, 61]) in the *Flavobacteriia*, was enriched by Tween80 0.01% and other Tween20 treatments, but most highly enriched by Tween20 0.005% (*P* value = 0.01) (**Figs. 6, S4**). Otu00210, 00216, 00306, and 00456 were enriched by at least two Tween20 treatments, with almost no enrichment by any Tween80 treatments (**Figs. 6, S4**). These OTUs included unclassified members of the *Verrucomicrobiaceae* and *Bacteroidetes,* both of which are consistently detected in other PA microbial communities [3, 8, 50–54, 57, 59, 61]. Otu00419, a *Synechococcus* representative, was correlated with the Tween20 treatments (**Fig. S4**) and enriched in Tween20 0.001% and 0.005% specifically (Figs. 6, S4). Although was also enriched by Tween80 0.005% (**Figs. 6, S4**), the effect was greater in the Tween20 0.005% treatment, indicating that the correlation of Otu00419 to Tween20 treatments was based on subtle differences. *Synechococcus* cyanobacteria are commonly detected in the PA fraction [8, 11, 58, 60, 62] and form aggregates in coastal waters, thus making them representative of the particulate organic matter pool [63].

Overall, our observed PA communities were dominated by bacteria; archaeal OTUs (nOTUs = 48) only constituted 0.048% ± 0.002% of the total (**OTU Table, FigShare**), however, two archaeal OTUs, Otu00739 and Otu00934, were still among the E-PA-OTUs (**Fig 6, S4**). Tween20 treatments generally performed poorly at dissociating archaeal OTUs, aside from Tween20 0.001% that enriched Otu00739. Of the Tween80 treatments, Tween80 0.005% performed best at enriching archaeal OTUs, nearly tripling the relative abundance of Otu00739 (*P* value = 0.02) and nearly doubling the relative abundance of Otu00934 (*P* value = 0.03) (**Fig. 6, S5**). Otu00934 and Otu00739 belonged to Marine Group II (99.6% identical), and Marine Group III (99.6% identical), respectively, via blastn alignment [64]. Both Marine Group II (MGII) and Marine Group III (MGIII) have been detected in PA fractions [65–67], further corroborating the significant increases in PA-OTUs that we are measuring.

We do not know the mechanism by which Tween80 outperformed Tween20 in enriching for the most abundant PA-OTUs, but we speculate that it stems from differences in the Tween20 and Tween80 side chains. Tween20 and Tween80 are both characterized as polysorbate nonionic detergents and they vary in their fatty acid side chains [68]: Tween20 has a saturated 12-carbon (12:0) lauric acid side chain whereas Tween80 has an unsaturated 18-carbon (18:1) oleic acid side chain [69]. In general, longer alkyl chains in fatty acids are more hydrophobic, and although double bonds in alkyl chains decrease hydrophobicity, oleic acid has almost twice the hydrophobicity value of lauric acid [70]. Thus the higher degree of hydrophobicity of Tween80 may better disrupt the microbial adhesion strategies, such as stalks or pili [45], which are typically hydrophobic in nature [71–73].

### Non-specific E-PA-OTUs

Some of the E-PA-OTUs were not associated with any particular Tween treatments, but rather increased across all treatments (**Figs. 4, 6, S3**). For example, Otu00333, an unclassified member of the *Verrucomicrobiaceae*, a group that is commonly detected in the PA fraction [8, 50–54], approximately tripled in relative abundance across all treatments (**Fig. 6**). Similarly, Otu00125 (unclassified *Flavobacteriaceae*), Otu00128 (*Crocinitiomix*), and Otu00326 (*Cryomorphaceae*), all members of *Bacteroidetes* that have been previously detected in PA communities [3, 8, 50, 56, 57, 59, 61], were also highly enriched across all treatments (**Fig. 6**). Otu00079, an unclassified member of *Proteobacteria*, and Otu00152, a *Phaeobacter* representative from within the *Alphaproteobacteria* (another group previously detected in the PA-fraction [9, 61, 74]), were evenly enriched across all Tween treatments as well. Otu00545, an unclassified member of the *Flavobacteriales*, was highly enriched across all treatments, averaging approximately 2-7-fold increases in relative abundance, except in Tween80 0.001%, where it increased approximately 17-fold (*P* value = 0.04) (**Fig. 6**). Despite being so highly enriched in Tween80 0.001%, Otu00545 was not included among the OTUs that significantly drove differences in PA-OTUs among Tween treatments (**Figs 4. S3**), likely due to overlapping fold changes in relative abundance between Tween20 and the other Tween80 treatments.

### Eukaryotic algal enrichment

The presence of chloroplasts among E-PA-OTUs (**Fig. 6**) was likely evidence of cell lysis as most eukaryotic phytoplankton will not pass through a 2.7µm pore-size filter. We identified unclassified chloroplast OTUs Otu00044, Otu00055, Otu00249 and Otu00304 as belonging to *Heterosigma akashiwo* (99.2% identical), *Fibrocapsa japonica* (100% identical), *Aureococcus anophagefferens* (99.6% identical), and an unknown marine eukaryote (99.21% identical), respectively. The first three are eukaryotic algae with cell sizes typically ranging 3-60µm [75–79]. We also observed diatom chloroplast OTUs, Otu00015 and Otu00514, that corresponded to *Skeletonema pseudocostatum* (98.8% identical), and *Bolidophyceae spp.* (99.6% identical), respectively. *Skeletonema pseudocostatum* is a chain-forming diatom with individual cell sizes ranging 2-9µm [80, 81] and *Bolidophyceae* are algae within the Stramenopiles, sister to diatoms, with cell sizes ranging 4-6µm [82]. These large cell sizes supported our hypothesis of Eukaryotic cell lysis and/or chain disruption by the Tween treatments. One might assume that higher concentrations of detergent would lead to more cell lysis, but instead we observed an overlap in eukaryotic chloroplast enrichment across the highest and lowest Tween concentrations (**Fig. 6**). Furthermore, our flow cytometry observations did not indicate that one treatment was causing increased cell mortality over another (**Figs. 1, S2**), even though the sample core on our flow cytometer was 16µm in diameter, and therefore possible for us to detect compromised larger cells with the live/dead stain. This suggests that Tween treatments do disrupt Eukaryotic cells, but the interactions are difficult to predict with the present data.

## Conclusion

Together, flow cytometry and community composition analyses indicated that Tween20 and Tween80 both effectively dissociate microbes from particles, but with different outcomes (**Figs. 2, 4, 6**). Tween80 treatments often dissociated a higher magnitude of PA cells, reaching upwards of a 70% increase in cell density (**Fig. 2**). Also, the cell mortality between Tween20 and Tween 80 treatments was consistently similar, indicating that Tween80 was just as gentle, but more effective than Tween20. Additionally, shaking at 185 RPM was a more reliable perturbation method compared with vortexing, particularly any longer than five minutes (**Fig. S2B**). At the OTU level, Tween80 demonstrated greater uniformity in the dissociated communities than Tween20 (**Fig. 4**), and significantly enriched the most abundant PA-OTUs, thus capturing the members that best represented the PA community. Tween80 treatments were also more effective in dissociating archaeal PA cells into the FL-fraction (**Fig. 6, S4**), particularly Tween80 0.005% (**Fig. 6**). We recommend sampling for the FL and PA communities separately, as the Tween treatment does affect the community composition of the FL community (**Fig. 3**). Tween80 also reliably and effectively dissociated a broad range of PA cells into the FL fraction across seasons and marine environments (**Figs. 2, 4, 5, S3, S5**). To limit the amount of detergent added to invaluable samples, capture a broad taxonomic representation of PA microbes with limited cell mortality, we recommend Tween80 0.005%, as it strikes a balance between a gentle dissociation without compromising dissociative power.

## Materials and Methods

### Environmental sample collection

For the 16S rRNA gene community composition analysis, we sampled 60 liters of seawater on November 20, 2021, from the Santa Monica Bay (33.99675°N, 118.48603°W) via manual surface water collection into four twenty-liter sterile polycarbonate carboys with five liters of headspace per carboy (Thermo Fisher Scientific, Waltham, MA). Sample processing began within one hour of sample collection. For flow cytometry analyses, subsequent sampling from the same Santa Monica Bay site occurred in 2023 on February 16 (winter), April 3 (spring), June 29 (summer), and November 26 (fall), with three liters of surface seawater collected into sterile four-liter carboys (Thermo Fisher Scientific, Waltham, MA) and sample processing beginning within one hour of sample collection. For additional flow cytometry analysis, samples were collected on May 15th, 2024 from the San Pedro Ocean Time-series (SPOT) sampling site (33.55°N, 118.4°W) to reflect microbial communities that are more characteristic of offshore waters [83] with offshore geochemistry [84]. Seven liters of surface seawater (2 m) was collected via Niskin bottles and transferred into a sterile 10L carboy (Thermo Fisher Scientific, Waltham, MA). Due to logistics of sample collection, the carboy was stored at close to ambient seawater temperature (20°C), and sample processing was performed within 18 hours of sample collection.

### Microbial dissociation from particles

We tested four concentrations of Tween20 (Sigma-Aldrich, St. Louis, MO) and Tween80 (Sigma-Aldrich, St. Louis, MO). We diluted stock solutions of 5% Tween20 and Tween80 (v/v) in sterile phosphate buffered saline (1x PBS) into surface seawater (sample volumes below) at final concentrations of 0.001%, 0.005%, 0.01%, and 0.1% (v/v), along with untreated controls. We separated particle-associated and free-living taxa via size-fractionation where cells caught on a 2.7 µm pore size GF/D filter (Whatman, Maidstone, United Kingdom) were considered particle-associated (PA) and cells that flowed -through the 2.7 µm filter were considered planktonic/free-living (FL). The logic of the experimental design was that PA cells that were previously collected on 2.7 µm filters would pass through these filters after treatment with the detergent and dissociation from particles. Thus, after dissociation, formerly PA cells would be detected in the 2.7 µm filtrate either via flow cytometry or collection on smaller filter pore sizes (0.22 µm). The relative success of the detergent would be quantified via the relative increase in total cells (evaluated via propidium iodide vs Sybr-green staining, see below) and/or PA community members in the 2.7 µm filtrate.

For flow cytometry analyses, we diluted Tween20 and Tween80 into sterile 50 mL falcon tubes (VWR, West Chester, PA) containing 30 mL surface seawater from the Santa Monica Bay and SPOT at the concentrations described above in triplicate. Samples were either vortexed at maximum speed for five or ten minutes, or they were shaken at 185 RPM for five or ten minutes. For vortexing at maximum speed, sample tubes were in a vertical/upright position on the vortexer. For 185 RPM shaking, sample tubes were secured in a horizontal position to increase the surface area for more mixing. After vortexing/shaking, we syringe-filtered the Tween-treated seawater through a sterile 2.7 µm 25 mm GF/D filter (Whatman, Maidstone, United Kingdom) and collected the filtrate in a new sterile 50 mL falcon tube. Each replicate received a new 50 mL sterile syringe (VWR, West Chester, PA) and sterile 25 mm GF/D filter. We processed control samples in the same manner; however, these samples received no Tween treatment and no physical perturbation. To understand the effect that the perturbation alone had on the microbial communities, we processed additional controls from the Santa Monica Bay winter and fall samples, along with the SPOT samples, without Tween but with both vortexing and shaking, separately, for five and ten minutes (**Figs. 2, S2**). We quantified cell concentrations from aliquots of the 2.7 µm filtrate for each treatment and replicate with a BD Accuri C6 Plus flow cytometer (BD, New Jersey, USA) after a 30-minute 1x SYBR-green (Lonza Bioscience, Basel, Switzerland) dark, room-temperature incubation. Separately, the number of dead cells was also quantified from aliquots of the 2.7 µm filtrate from each treatment and replicate using the same flow cytometer after a 30-minute 1x propidium iodide (Thermo Fisher Scientific, Waltham, MA) dark, room-temperature incubation. Results were visualized in R v.4.4.0 [85, 86] using ggplot2 [87].

For 16S rRNA gene community composition analysis, we diluted the four different concentrations of Tween20 and Tween80 into separate, sterilized 10 L carboys containing 7 L of surface seawater from the Santa Monica Bay in triplicate that we shook manually for 5 minutes. The free-living (Sterivex) Tween20 0.1% only received duplicate samples. Subsequently, 2 L of the Tween-treated seawater were filtered via peristaltic pumping (Cole Parmer 77601-10, Vernon Hills, IL) with an in-line (Masterflex I/P tubing, Germany) sterile 2.7 µm 47mm GF/D filter (Whatman, Maidstone, United Kingdom) and a sterile 0.22 µm Sterivex filter (Sigma-Aldrich, St. Louis, MO). Each replicate received new sterile filters, and filter lines were flushed with 1 L of 70% ethanol and 2 L MilliQ water between samples. Control samples that received no physical perturbation and no Tween treatment were also filtered in triplicate through the same in-line filtration setup. Prior to filtration, filter lines were sterilized with equal volumes of 0.1N HCl (Sigma-Aldrich, St. Louis, MO), then 0.1N NaOH (Sigma-Aldrich, St. Louis, MO), then MilliQ water. After filtration, samples were immediately stored at −20°C until further processing.

### DNA extraction and sequencing

We extracted DNA for 16S rRNA gene sequencing from all GF/D and Sterivex filters across all treatments based on a phenol-chloroform-isoamyl alcohol extraction method and the Griffiths Method [88]. Briefly, we removed filters from −20°C storage and thawed them on ice. We tore GF/D filters into quarters using sterile forceps and placed each quarter into a sterile microcentrifuge tube. We sliced Sterivex filters in half using sterile razor blades and placed each half into a sterile microcentrifuge tube with sterile forceps. Chemical lysis was performed by adding 557 µL 1x Tris-EDTA buffer (Sigma-Aldrich, St. Louis, MO), 139 µL 10% SDS (Sigma-Aldrich, St. Louis, MO), and 35 µL lysozyme (1.4mg/mL final concentration, Sigma-Aldrich, St. Louis, MO) to each tube and vortexing for 30 seconds, then incubating at 37°C for 30 minutes. After incubation, we added 70 µL of proteinase K (0.7 mg/mL final concentration, Sigma-Aldrich, St. Louis, MO) to each tube, vortexed for 20 seconds, then incubated overnight at 55°C [89]. After overnight incubation, the we separated the lysate from the GF/D filters by transferring the GF/D filter + lysate into sterile 0.45 µm pore size Costar Spin-X tubes (CLS8162, Sigma-Aldrich, St. Louis, MO), and centrifuged at 12,000 x g for 1 minute, then the lysate from the GF/D filter was transferred into a new sterile microcentrifuge tube. Lysate separation for the Sterivex filter was unnecessary as the polyethersulfone membrane is dissolved during extraction. After overnight lysis incubation and lysate separation from GF/D filters, we added 500 µL CTAB extraction buffer [88] and 500 µL phenol-chloroform-isoamyl alcohol (25:24:1, pH 8.0, Sigma-Aldrich, St. Louis, MO) to the lysate, vortexed for 30 seconds, then centrifuged at 16,000 x g at 4°C for five minutes. We transferred the aqueous layer to a new sterile microcentrifuge tube and equal volume of chloroform-isoamyl alcohol (24:1, Sigma-Aldrich, St. Louis, MO) was added then centrifuged again at 16,000 x g at 4°C for five minutes. We again transferred the aqueous layer to a new sterile microcentrifuge tube and precipitated the DNA for one hour at room temperature with a 0.1 x aqueous volume 3M sodium acetate (pH 5.2, Sigma-Aldrich, St. Louis, MO), and 0.6 x aqueous volume isopropanol [90], then centrifuged at 16,000 x g at 4°C for fifteen minutes. We washed the DNA pellet with 150uL of ice-cold 100% (v/v) molecular grade ethanol (Sigma-Aldrich, St. Louis, MO) and centrifuged again at 16,000 x g at 4°C for fifteen minutes, air dried, and resuspended in 50 µL of molecular grade water. Pure DNA was achieved by incubating resuspended DNA with RNase A (Promega, Madison, WI) for 10 minutes according to manufacturer’s instructions. Purified DNA was stored at −80°C until sequencing.

Prokaryotic genomic DNA was amplified at the V4 region of the 16S rRNA gene using the 515F and the modified 806R primer set [91, 92], and the QuantaBio’s AccuStart II PCR ToughMix (QuantaBio, Beverly. MA). Amplicons were sequenced at Argonne National Laboratory on an Illumina MiSeq using paired-end 250 base-pair reads using the v2.5 TruSeq Paired End MiSeq flow cell/cluster kit (Illumina Inc., San Diego, CA) according to manufacturer’s instructions.[91]

### Community composition

To assess how prokaryotic community composition changed in response to Tween treatments, raw 16S rRNA gene sequences were analyzed using Mothur v.1.48.0 [93] and the Silva v.132 database [94], keeping replicates separate. We first assembled 16S rRNA gene amplicon sequences into contigs, and discarded contigs with ambiguous bases, nucleotide repeats greater than 8 base-pairs (bp), or those that were greater than 275 bp in length. We aligned and classified contigs via the Silva v.132 database, and removed chimeric contigs, with Mothur v.1.48.0. Contigs that were classified as “unknown” were removed, and the remaining contigs were clustered into Operational Taxonomic Units (OTU) with a 0.03 dissimilarity threshold resulting in 30,258 OTUs with a mean length of 252bp.

To assess the overall dissimilarity in OTU composition between size fractions and Tween treatments, we calculated Bray-Curtis dissimilarity [95] distance matrices and applied them toward Principal Coordinate Analysis [96] (PCoA) in Mothur v.1.48.0 [87]. We quantified the distribution of OTUs across technical replicates for all treatments in R v.4.4.0 using VennDiagram [97]. A PA-OTU was defined as such if it was present across all three GF/D filters, and vice versa for the FL-OTUs from control Sterivex filter replicates. To determine if PA-OTUs were becoming enriched in the FL fraction after Tween treatment, first we calculated fold changes in the relative abundance (RA) of PA-OTUs across all Tween treatments in the FL fraction. A PA-OTU with a fold-change in relative abundance greater than one in the FL-fraction was considered to have increased. We also defined whether an increased PA-OTU was “abundant” if it had a relative abundance of ≥ 0.1% in the control PA community; “rare” increased PA-OTUs were those with < 0.1% relative abundance in the control PA community. Second, focusing on the PA-OTUs with more than 1-fold increased abundance, we calculated Bray-Curtis dissimilarity [95] distance matrices and performed a PCoA [96] on this group (nOTU=316). Third, we fitted vectors for significantly increased PA-OTUs onto the ordination using the “envfit” function from the Vegan [98] package. Fourth, we then used the Wilcoxon signed-rank test [99] in R v4.4.0 [85, 86] to evaluate which of the increased PA-OTUs were significantly enriched in each of the Tween treatments relative to controls (E-PA-OTUs) and visualized those results in R v.4.4.0 [85, 86] using gglot2 [87]. If an OTU within the E-PA-OTUs was classified as “unclassified bacteria” we attempted to improve the taxonomic classification manually by aligning the OTU consensus sequence (**Representative OTU sequences fasta, FigShare**) to the NCBI *nr* database [100] via BLASTN [64].

## Data Availability

Raw sequencing fastq files are available on NCBI under BioProject PRJNA1134785. Accessory data and scripts used for 16S rRNA gene analyses, statistical tests, and data visualization, are available on FigShare (https://figshare.com/account/projects/256418/articles/29565185).

## Acknowledgements

The authors thank Shelby J. Barnes and Brittany Bennet for their helpful advice and suggestions during the optimization and development of the experimental protocol. Additionally, we thank Zachary Henning, Hasti Asrari, Troy Gunderson, and the crew of the *R/V Yellowfin* for collecting seawater from the San Pedro Ocean Time Series study site. Funding for this research was provided by a National Science Foundation grant (OCE-1945279) to J.C.T.

## Supplemental Figure Captions

**Figure S1. Additional flow cytometry data across seasons.** X-axes indicate treatments. The left y-axis represents the cell density determined by the SYBR-Green fluorescent stain (black text). The right y-axis represents the percentage of dead cells within the sample as determined by the Propidium Iodide fluorescent stain (grey text). The percentages above the boxplots indicate the average percent increase in cell density, or percent cell death, relative to the control. The boxplots describe the distribution of the data, with the boxes indicating the lower and upper quartiles, the horizontal line indicating the median, and the whiskers showing the minimum and maximum values. A) Samples collected from the Santa Monica Bay on June 29, 2023 (representing the summer season). Samples were either shaken at 185 RPM or vortexed, both for 10 minutes. The percentages above the box plots are derived from the control that also received either the shaking or vortexing perturbation treatment. B) Samples collected from the Santa Monica Bay on November 26, 2023 (representing the fall season). Samples were vortexed for either 5 minutes or 10 minutes. The percentages above the box plots are derived from the control that also received the vortexing perturbation treatment.

**Figure S2. Percentage of OTUs distributed across replicates.** X-axis indicates the OTU distribution category; y-axis indicates the percentage of OTUs in each category. The boxplots describe the distribution of the data, with the boxes indicating the lower and upper quartiles, the horizontal line indicating the median, and the whiskers showing the minimum and maximum values. The color of the boxplot corresponds to the control or experimental treatment, according to the key. Tween20 0.1% sterivex filters only received duplicate rather than triplicate replicates (*).

**Figure S3. Principal Coordinate Analysis of the particle-associated communities that increased in the free-living fraction in response to Tween treatments (*P* value < 0.05)**. The colors of the particle-associated (▴) shapes correspond to the Tween treatment, as indicated by the key. The crosses indicate the centroid points for each treatment. Vectors represent OTUs with significant relationships (*P* value < 0.05) with ordination axes. Red vector labels represent an OTU that significantly increased in the FL fraction in response to at least one Tween treatment (Figure 6, Table S3).

**Figure S4. E-PA-OTUs with fold change increases less than 3.5.** Same data as in Figure 6, subsetted for E-PA-OTUs with less enrichment to improve resolution. The E-PA-OTUs that significantly increased in at least one Tween treatment (Table S3) are organized by OTU number (most overall relative abundance to least). The x-axis indicates the fold change in relative abundance, and the y-axis shows OTUs with their taxonomy. The colors of the data points correspond to the Tween treatment, as indicated by the key. Datapoints and error bars represent the mean and variation in fold-change in relative abundance across triplicates.

**Figure S5. E-PA-OTU enrichment by Tween treatment, including Tween20 0.1%.** The E-PA-OTUs that significantly increased in at least one Tween treatment (Table S3) are organized by OTU number (most overall relative abundance to least). The x-axis indicates the fold change in relative abundance, and the y-axis shows OTUs with their taxonomy. The colors of the data points correspond to the Tween treatment, as indicated by the key. Datapoints and error bars represent the mean and variation in fold-change in relative abundance across triplicates, except for Tween20 0.1%, which come from duplicates.

## Supplemental Table Captions

**Table S1. PA-OTUs that increased with fold-change greater than one in response to Tween treatments.** Relative abundance for each OTU is displayed in both the Tween treatments and controls, along with the change in relative abundance in the Tween treatment relative to the control, and the corresponding fold change in relative abundance for each OTU.

**Table S2. Classification of increased PA-OTUs as either abundant or rare.** Relative abundance, taxonomy, and raw count data for each OTU derived from the control PA treatment. An OTU was considered abundant if it had a relative abundance greater than 0.1%.

**Table S3. Statistical enrichment of PA-OTUs.** Wilcoxon statistical significance of E-PA-OTUs for each Tween treatment, *P* values < 0.05.

