## Supplementary figures and images for "Detergent-based separation of microbes from marine particles"

### Figure S1

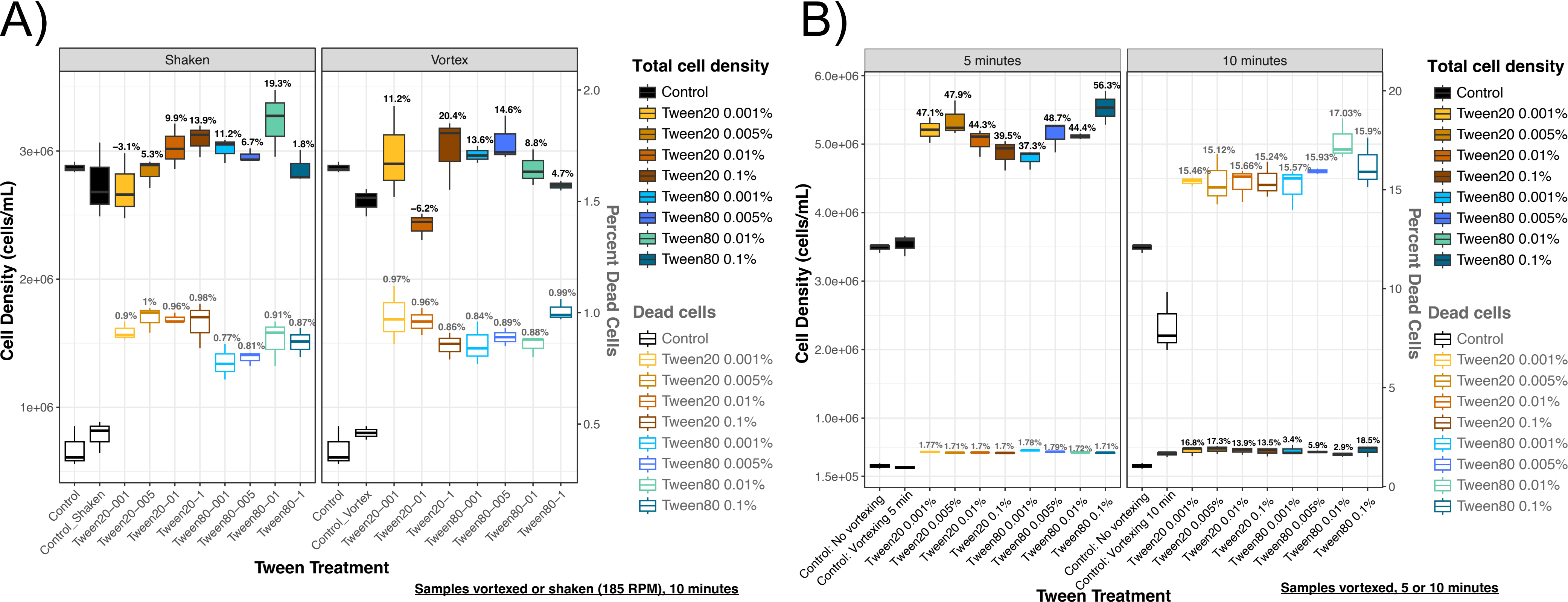

### Figure S2

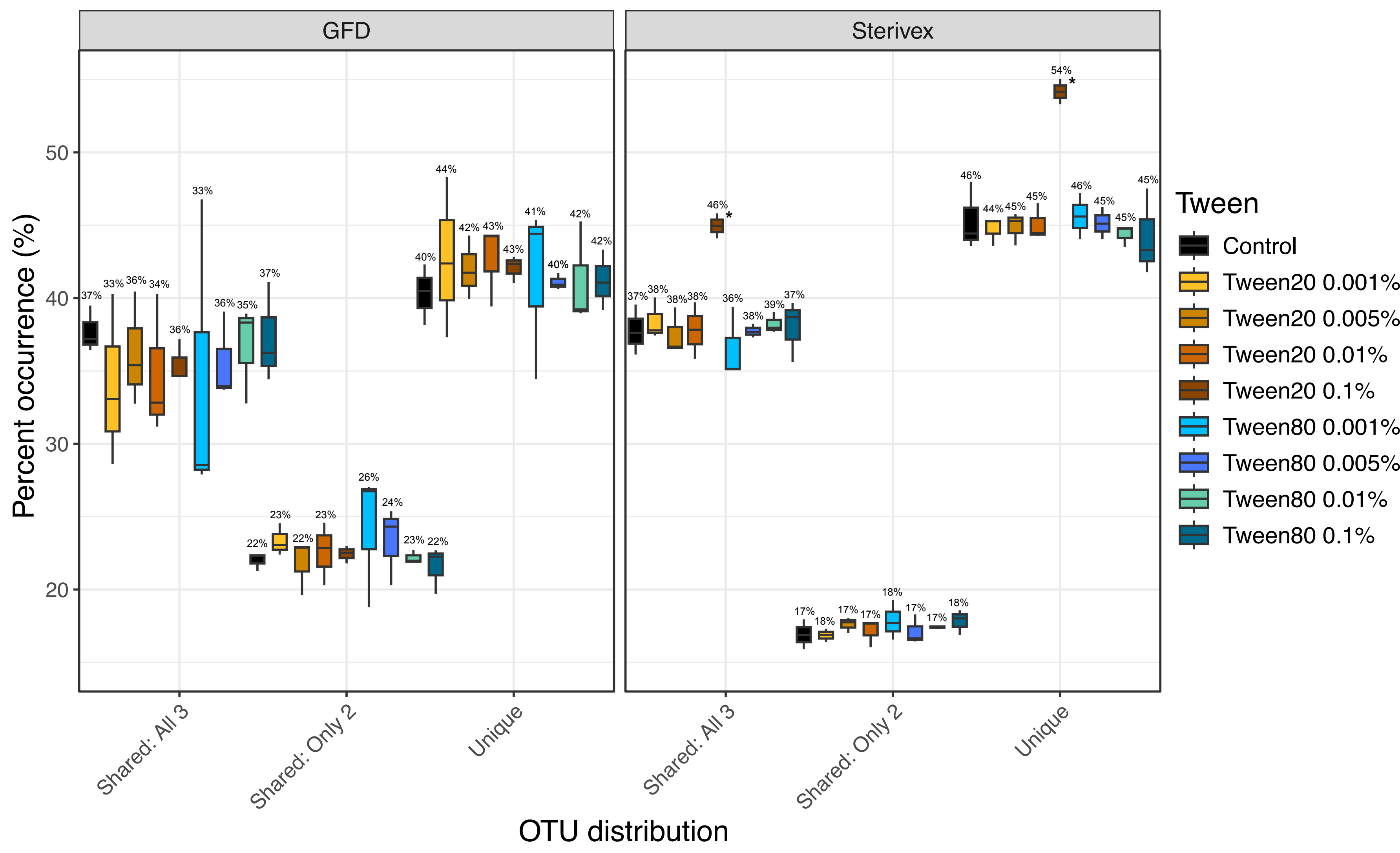

### Figure S3

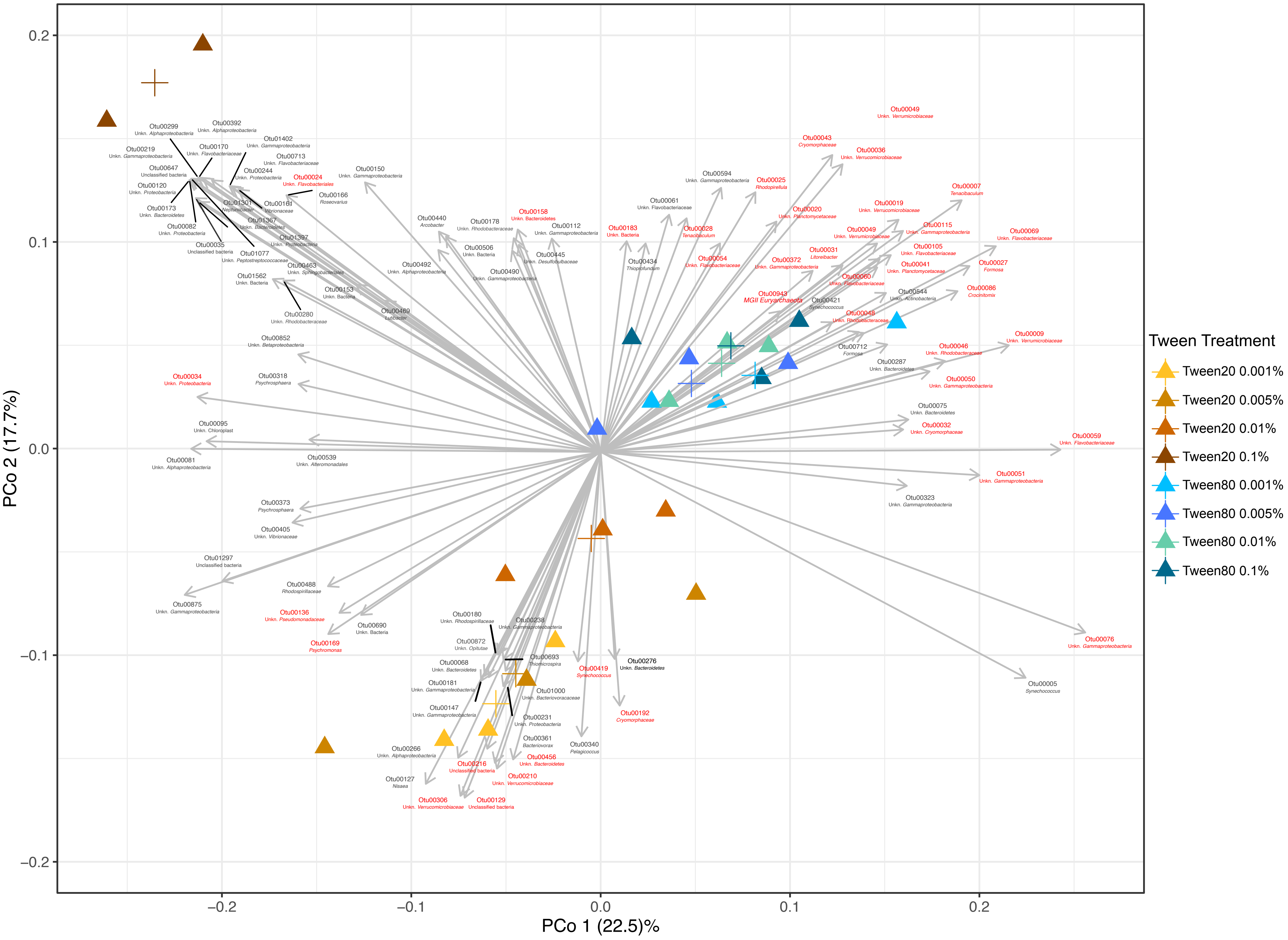

### Figure S4

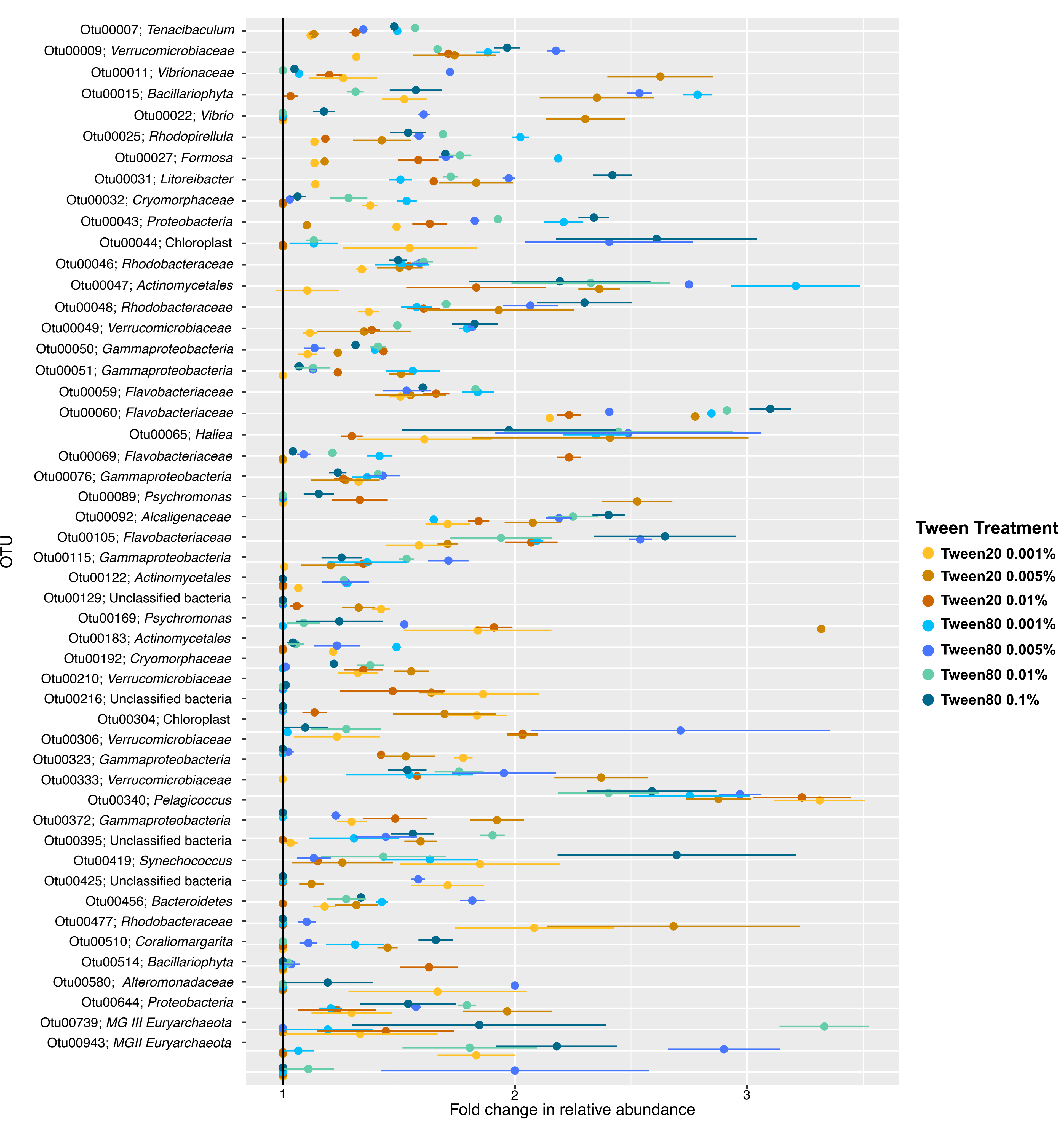

### Figure S5

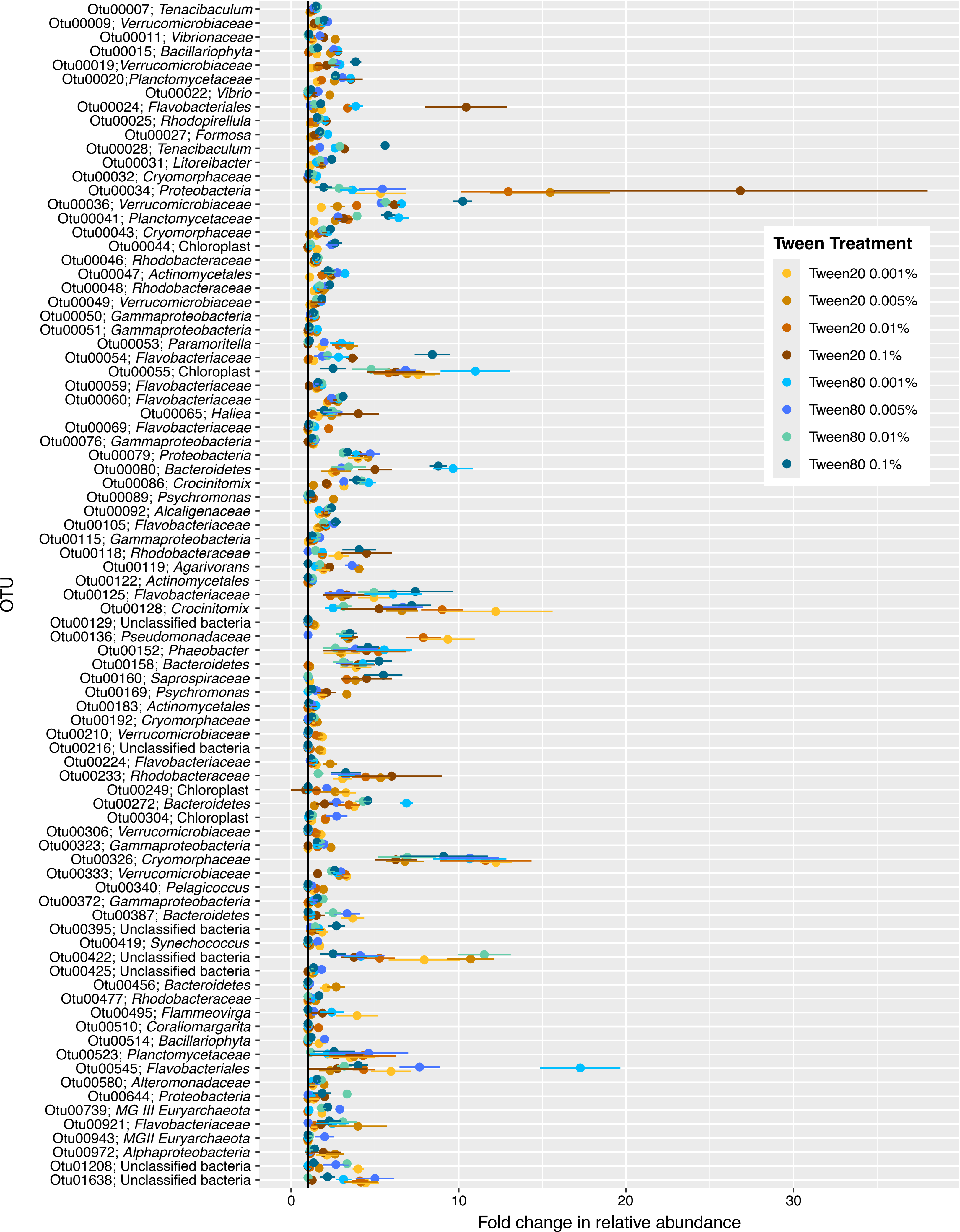
